# Aggressive nitrogen assimilation during exponential growth in *Chlamydomonas reinhardtii*

**DOI:** 10.1101/2025.11.02.686124

**Authors:** Miku Tibule, Windula Mallawarachchi, Oyinkansola Oyateru, Mahtab Ziaeian, Minsung Kim, Hasni Nimalka Dharmasiri, Jae-Hyeok Lee

**Affiliations:** Department of Biological Sciences, University of Manitoba, Winnipeg, Manitoba R3T 2N2, Canada

**Keywords:** nitrogen use efficiency, *Chlamydomonas reinhardtii*, batch culture, ammonium, nitrate, urea, intracellular nitrogen storage

## Abstract

Bioavailable nitrogen (N) is a key limiting factor for global biomass production. In dynamic environments, organisms must employ effective nitrogen use strategies (NUS) to capitalize on episodic N availability. The recent discovery of intracellular guanine crystals in diverse algal species has prompted questions about their roles in NUS. To investigate NUS in *Chlamydomonas reinhardtii,* a model photosynthetic eukaryote, we compared N-to-biomass conversion rate of mixotrophic batch cultures fed with three common N sources, ammonium, nitrate, and urea. Saturating growth was achieved at 4 mM ammonium, 4 mM nitrate, and 2 mM urea, indicating comparable molar N utilization efficiency. Residual N measurements revealed that approximately 1.2 unit of optical density (O.D. at 680 nm) biomass was produced per mM N under sub-saturating conditions, while biomass accumulation per N decreased at above-saturating conditions. To estimate N storage capacity, we tracked N uptake kinetics in high-density culture following an N pulse, showing rapid assimilation of 3∼6 mM N per O.D. biomass. N source-specific transcriptome revealed N source-specific regulation of assimilation pathways and transporter genes in support of effective NUS in *C. reinhardtii*. These findings support a model in which *C. reinhardtii* can rapidly acquire and store N during exponential growth, enabling sustained growth and metabolism during periods of N scarcity. Observed variation in N storage capacity across growth stages and N conditions predicts the regulatory mechanisms governing N partitioning and storage. This study highlights flexible NUS in microalgae, offering insights for improving N assimilation capacity and resilience in agricultural and aquacultural crops.

## 1. Introduction

Nitrogen (N) is an essential macronutrient and a major limiting factor in biomass production across ecosystems. Although abundant in the atmosphere, bioavailable N forms are historically finite, making nitrogen use efficiency (NUE) a central focus of agricultural and ecological research (Kopittke et al., 2019; Bailey-Serres et al., 2019). The industrial synthesis of ammonia via the Haber-Bosch process disrupted the natural N cycle, unlocking the potential of autotrophic growth in crops that sustain the nutritional needs of the expanding human population (Erisman et al., 2008; Good et al., 2011).

However, synthetic N fertilizers now account for roughly one-third of the 10% agricultural contribution to global greenhouse emissions, and an estimated 50% of applied N is lost to leaching, leading to eutrophication and nutrient imbalance in ecosystems (Govindasamy et al., 2023).

Photosynthetic organisms can exploit multiple N sources available in aquatic and soil environments, such as ammonium (NH_4_^+^), nitrate (NO_3_^-^), and urea (Nunes-Nesi et al, 2010; Calatrava et al., 2023). These N sources enter cells through low-affinity nonselective channels or high-affinity selective transporters and must ultimately be converted to ammonium before assimilation into amino acids (Nunes-Nesi et al., 2010; Krapp, 2015). The subsequent incorporation of NH_4_^+^ into glutamine and glutamate, primarily catalyzed by the glutamine synthetase/glutamate synthase (GS/GOGAT) cycle, defines the key step of N assimilation and thereby its efficiency determines N-dependent growth kinetics (Peltier and Thibault, 1983; Suzuki and Knaff, 2005; Thomsen et al., 2014). Continuous operation of the GS/GOGAT cycle requires a steady supply of carbon (C) skeletons in the form of 2-oxoglutarate (2-OG), underscoring the central importance of coordinated C and N metabolism for sustainable growth (Nunes-Nesi et al., 2010; Kramer and Evans, 2010).

Agricultural systems optimized for yield experience chronic N excess, which not only promote NO_3_^-^ leaching but can also inhibit photosynthesis (Hachiya and Sakakibara, 2016). High concentrations of NH_4_^+^ are known to inhibit growth in higher plants and microalgae by disrupting the C/N balance and suppressing photosynthesis (Hachiya and Sakakibara, 2016). A recent genetic study showed that disruption of chloroplastic GS eliminates ammonium-induced growth inhibition (Hachiya et al., 2021), suggesting that excess NH_4_^+^ imposes a metabolic imbalance that feeds back to suppress photosynthetic activity. Interestingly, such inhibition is not observed under NO_3_^-^ excess, implying that cells differentially sense and respond to distinct N sources.

In plants, NO_3_^-^ acts as both a nutrient and a signal activating two sensor systems: the dual-affinity transporter CHL1/NRT1.1, which mediates calcium-dependent signaling at the plasma membrane, and the RWP-RK family transcriptional regulator NLP7, which translocates to the nucleus upon nitrate binding to activate target genes (Ho et al., 2009; Xu et al., 2016; Liu et al., 2022). In *Chlamydomonas reinhardtii*, the orthologous transcriptional regulator NIT2 plays an analogous ancestral role, though the molecular details of NO_3_^-^ signaling remain incompletely understood (Schnell and Lefebvre, 1993; Carmargo et al., 2007). By contrast, NH_4_^+^-dependent signaling mechanisms in plants are still poorly defined (Britto et al., 2002; Ying and von Wiren, 2018; Liu et al., 2021), likely reflecting the relatively minor contribution of NH_4_^+^ to plant NUS compare with nitrate in the pre-anthropogenic earth.

Typical plant NUE studies assess growth responses over days or weeks under different N sources and concentrations (Masclaux-Daubresse et al., 2010; Chen et al., 2020). However, because the majority of plant biomass is composed of carbohydrates, NUE in plants often reflects conditions that best sustain photosynthetic efficiency rather than N assimilation per se. Given that N assimilation also critically depends on organic C supply, disentangling N-dependent growth responses from C-related effects is inherently challenging in obligate photoautotrophs (Cheng et al., 2023).

Unicellular green algae such as *C. reinhardtii* offer a powerful model for dissecting the molecular basis of photosynthetic and nutrient assimilation processes (Saroussi et al., 2017; Strenkert et al., 2019; Fauser et al., 2022). *Chlamydomonas* can grow heterotrophically in the absence of photosynthesis, while its photoautotrophic metabolism closely parallels that of plants (Catalanotti et al., 2013; Vuong et al., 2015). This dual metabolic flexibility, coupled with experimental tractability through nutrient manipulation, metabolite/inhibitor supplementation, and genetic perturbation, enables rapid and quantitative analysis of nutrient responses (Johnson and Alric, 2012; Bogaert et al., 2019). Moreover, as a unicellular organism lacking tissue compartmentalization and central vacuole as a potential nutrient reservoir, *Chlamydomonas* likely employs more elaborate N use strategies than multicellular plants. Its pyrenoid-based carbon-concentrating mechanism, a specialized adaptation for efficient CO_2_ fixation, exemplifies algal innovations absent in plants (He et al., 2023). Therefore, understanding and integrating these algae-based NUS into plant systems could inspire new strategies for improving agricultural nutrient efficiency and productivity (Atkinson et al, 2016).

As the first step toward establishing *Chlamydomonas* as a model for NUE research, we implemented a strict quantitative protocol measuring biomass accumulation per unit N consumed. This approach generated a reproducible N economy profile of *Chlamydomonas* culture, serving as a standard for future studies. Our findings reveal highly potent N storage mechanisms that accumulate N in non-biomass forms, reaching over 100% of existing biomass under excess N conditions. These data are consistent with the observed accumulation of intracellular guanine crystals, suggesting that *Chlamydomonas* possesses an extensive capacity for regulated N storage beyond immediate growth needs.

## 2. Materials and Methods

### 2.1. Strains and culture conditions

*C. reinhardtii* wild-type and mutant strains were obtained from the Chlamydomonas Resource Center (https://www.chlamycollection.org). The wild-type strains included CC-5370 (mt–), CC-5119 (mt–), and CC-5082 (mt+). All strains were maintained on tris-acetate-phosphate (TAP) agar medium containing 8.0 mM NH_4_Cl at pH 7.0 (Gorman and Levine, 1965). Liquid cultures were grown under mixotrophic conditions (17 mM acetate) at 23 °C under continuous illumination (100 µmol photons m⁻² s⁻¹) with shaking (125 rpm) or aeration in a multicapacitor photobioreactor. For experimental treatments, N sources (NH_4_Cl KNO_3_, or urea) were supplied at concentrations ranging from 0.5 to 16 mM. Nitrogen-free TAP (NF-TAP) medium was prepared by omitting all N components.

### 2.2. Growth monitoring and biomass estimation

Growth kinetics of CC-5370 and CC-5119 were monitored in a Multi-Cultivator MC 1000-OD (Photon Systems Instruments) with a working volume of 80 mL. The cultures were maintained at 23 °C and illuminated with white LEDs at 100 µmol photons m^-2^ s^-1^. Optical densities at 680 nm and 720 nm were automatically recorded at 5 min intervals. Each experiment was performed with duplicates and independently repeated at least three times for reproducibility. Culture vessels were randomly positioned in the reactor to minimize sensor bias. Since CC-5370 and CC-5119 share the 21gr (mt–) background, their growth kinetics were comparable in ammonium- and nitrate-fed cultures and were combined for analysis. Distinct urea-fed growth patterns were analyzed separately.

### 2.3. Optical density as a biomass proxy

Optical density at 680 nm (OD_680_) was used as a proxy for biomass instead of cell counts, because *C. reinhardtii* cell size varies more than tenfold through the cell cycle. Under our conditions, 1 OD_680_ corresponds to approximately 5 × 10^6^–1 × 10^7^ cells mL^-1^ (or 4 × 10^8^–8 × 10^8^ cells per 80 mL culture). As OD_680_ primarily reflects chlorophyll content, which can vary with physiological state, the OD_680_/OD_720_ ratio was monitored as an indicator of chlorophyll integrity. Healthy mixotrophic cultures maintained a ratio of 2.5–3.0, whereas urea-fed cultures at ≤ 1 mM exhibited a ratio of 1.5–2.0 and turned yellow at stationary phase.

### 2.4. Growth curve analysis

Raw OD data were screened for anomalies (e.g., spikes or baseline shifts caused by sensor obstruction or drift). Data were smoothed by sigmoidal fitting:

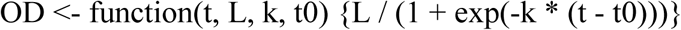

where *L* = maximum OD (asymptote), *k* = growth rate constant, *t0* = time at the inflection point. Growth rates were expressed as doubling rates per day (Δlog2OD per 24 h). All analyses were conducted using a custom Python script developed with Grok3-AI and Google Colab, available at https://github.com/Chlamy18/GrowthCurveAnalysis.

### 2.5. Culture harvesting

Samples (5 mL) were collected at lag, mid-log, and stationary phases. Cell counts were performed using a haemocytometer after fixation in 2% glutaraldehyde. The supernatants were stored at 4 °C for subsequent N quantification. For N storage assays, triplicate cultures of CC-5370 and CC-5119 were grown in 8 mM N medium. Upon reaching stationary phase, each culture was diluted 1:1 with fresh 8 mM N medium (N pulse). Samples were taken 10 h after NH_4_^+^ addition and 24 h after nitrate addition, reflecting the slower nitrate assimilation rate. N remobilization was assessed by resuspending mid-log or stationary-phase cells in NF-TAP and monitoring changes in OD_680_.

### 2.6. Nitrogen quantification

For ammonium quantitation, each sample was mixed with 0.1 vol. Nessler reagent (HI93715, Hanna Instruments), incubated for 10 min at room temperature, and measured at 410 nm. Calibration was performed with NH_4_Cl standards (0.05–1.5 mM) prepared in NF-TAP. Nitrate was quantified by the perchloric acid method (Cawse, 1967). Samples were treated with 0.5% (v/v) HClO_4_ and measured at 210 nm using a NanoDrop One spectrophotometer (Thermo Fisher Scientific). Standard curves were generated with KNO_3_ solutions (0.025–3 mM).

Urea was quantified by the Nessler method following urease-mediated hydrolysis into ammonium. Samples (125 µL) were incubated with an equal volume of urease solution (5 U; U1875, Sigma-Millipore) in 50 mM phosphate buffer for 1 h at 30 °C, and released ammonium was determined by the Nessler assay. Urea standards (0.025–2 mM) were processed in parallel. Control assays without urease confirmed negligible spontaneous urea degradation.

### 2.7. Fluorescence microscopy

To visualize nutrient deposits inside the cells at stationary phase, cells were laid on 0.5% low melting agarose without fixation and observed using a Zeiss LSM 700 laser confocal microscope with a 20x air and 40x oil-emersion objective lens. Guanine crystals were visualized by reflection imaging using 488 nm excitation and 479–489 nm emission according to Mojzes et al. (2020). Polyphosphate granules were stained with 10 µM DAPI (from 20 mM stock in water) and imaged using 405 nm excitation and 525– 550 nm emission following the protocol by Liang et al. (2025).

### 2.8. Gene expression analysis

Expression data for N assimilation and transporter genes were compiled from published and in-house RNA-seq datasets: (i) urea vs. ammonium transcriptome from Liang (2025; CC-125 *nit2* mt+), (ii) nitrate vs. ammonium transcriptome from this study (CC-5370 *NIT2* mt–), and (iii) ammonium vs. N-starvation transcriptome from Lee et al. (2025; CC-125-derived *nit2* mt+). All datasets were generated under TAP-based mixotrophic conditions. Sequencing and read mapping was done following the protocol by Lee et al. (2025). In brief, sequencing reads (15–20 million paired-end per sample) were mapped to the *C. reinhardtii* v6.1 genome using STAR aligner (v2.7; Dobin et al., 2013), and expression levels were estimated as fragments per kilobase per million reads (FPKM) with StringTie (v2.0; Pertea et al., 2014). Reported FPKM values represent means of two or three biological replicates.

### 2.9. Statistical analysis

All experiments were conducted with at least three biological replicates unless otherwise stated. Growth rate and nitrogen concentration data are presented as means ± standard errors (SE).

## 3. Results

### 3.1. Growth Kinetics and Saturation Points Vary by N Source

Numerous studies have examined how nutrient conditions affect biomass production (Freudenberg, 2021; Kropat et al., 2011; Lauersen, 2016; Bogaert et al., 2019). However, the stoichiometric relationship between nutrient availability and biomass yield has not been systemically assessed. To define the N stoichiometry underlying *Chlamydomonas* biomass production, we monitored OD_680_-based biomass accumulation across a range of N concentrations, from limiting to excess, under mixotrophic conditions, where growth typically saturate due to depletion of either N or acetate (Bogaeart et al., 2019).

Biomass production saturated at 4 mM NH_4_^+^, 4 mM NO_3_^-^, and 2 mM urea (Figures 1–3). Among these, NH_4_^+^ supported the fastest growth, achieving up to four doublings per day, whereas NO_3_^-^- and urea-fed cultures reached maximum rates of approximately three doublings per day. We defined a growth rate of 0.5 doublings per day as the threshold for entry into stationary phase (red dashed line in panels B in Figures 1-3).

**Fig. 1.**
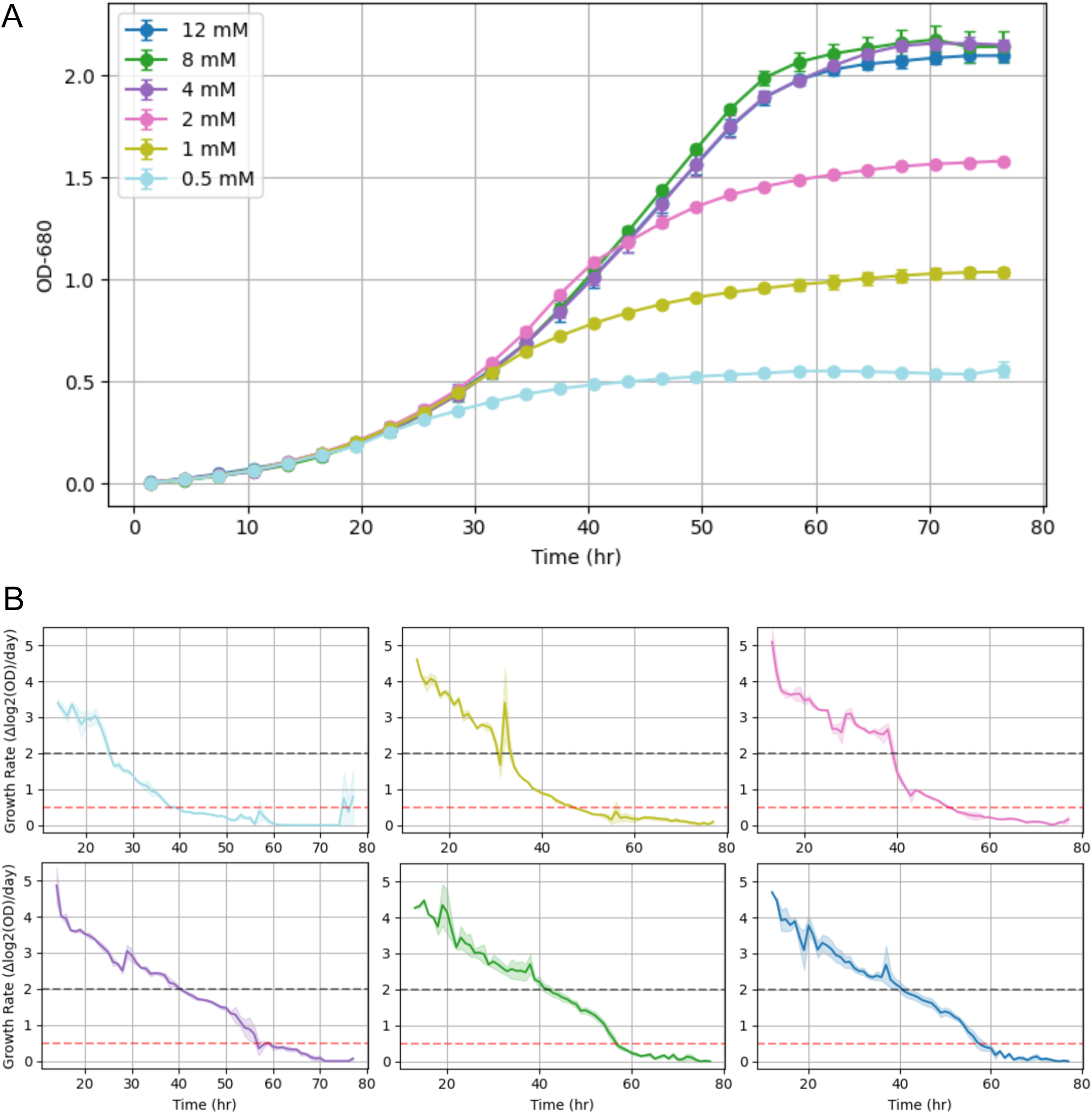
Ammonium concentration–dependent growth kinetics and doubling rates. (A) Growth kinetics of *C. reinhardtii* cultured under varying initial concentrations of ammonium chloride (0.5–12 mM) in mixotrophic TAP medium. Biomass accumulation was monitored by OD_680_ over time. (B) Growth rate (expressed as doublings per day) in one-hour sliding windows was calculated based on OD_680_ measurements and moving averages in one-hour sliding windows were plotted over time for each N concentration at OD_680_ ≥ 0.1. Black dashed line indicates 2.0 as a threshold for N-replete exponential growth, and red dashed line indicates 0.5 as a threshold for the entry into stationary phase.

**Fig. 2.**
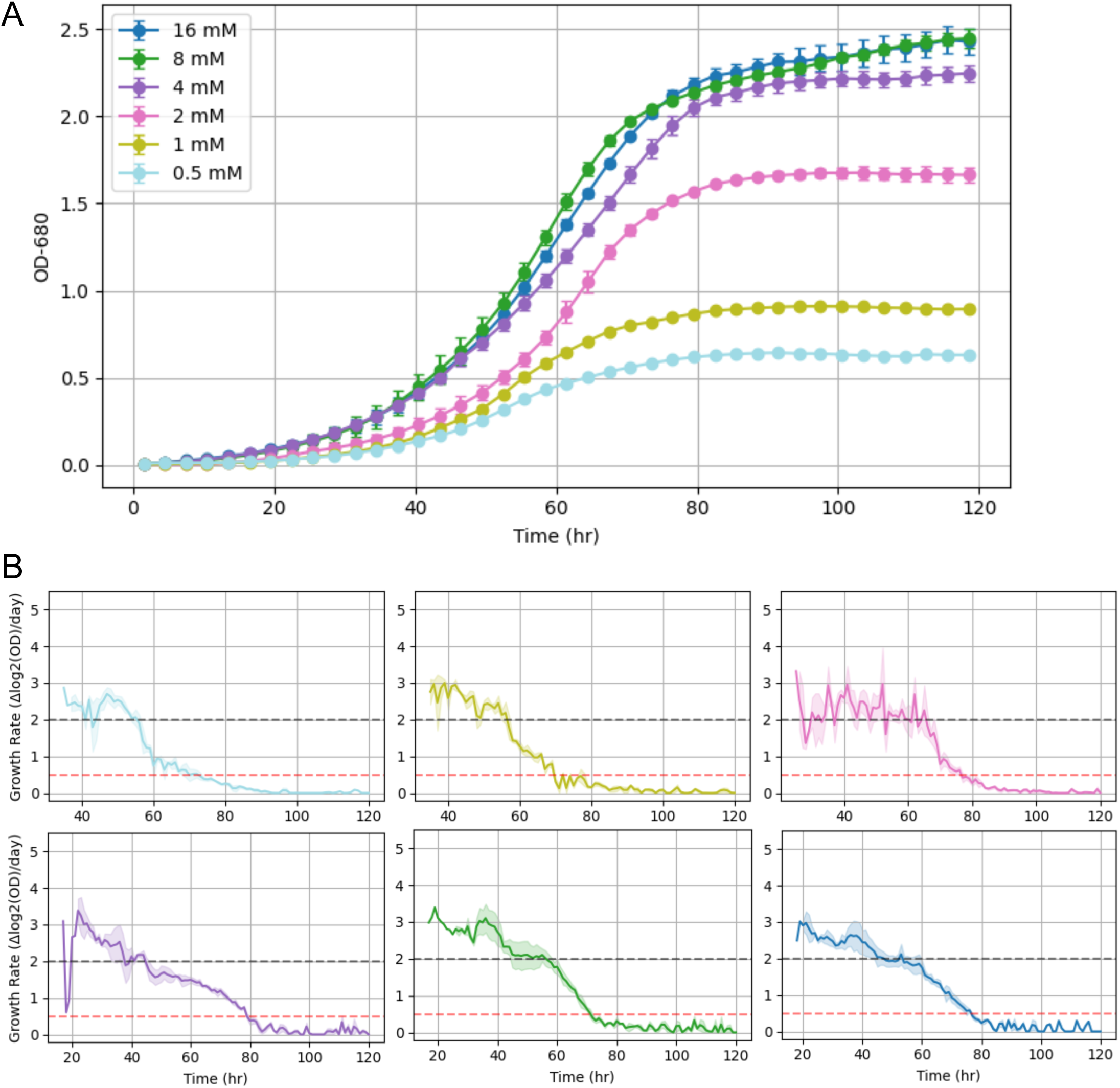
Nitrate-dependent growth dynamics and doubling rates. (A) Growth curves of *C. reinhardtii* cultured with varying initial concentrations of potassium nitrate (0.5–16 mM) under mixotrophic conditions. Biomass accumulation was monitored by OD_680_ over time. (B) Growth rate (expressed as doublings per day) in one-hour sliding windows was calculated based on OD_680_ measurements and moving averages in one-hour sliding windows were plotted over time for each N concentration at OD_680_ ≥ 0.1. Black dashed line indicates 2.0 as a threshold for N-replete exponential growth, and red dashed line indicates 0.5 as a threshold for the entry into stationary phase.

**Fig. 3.**
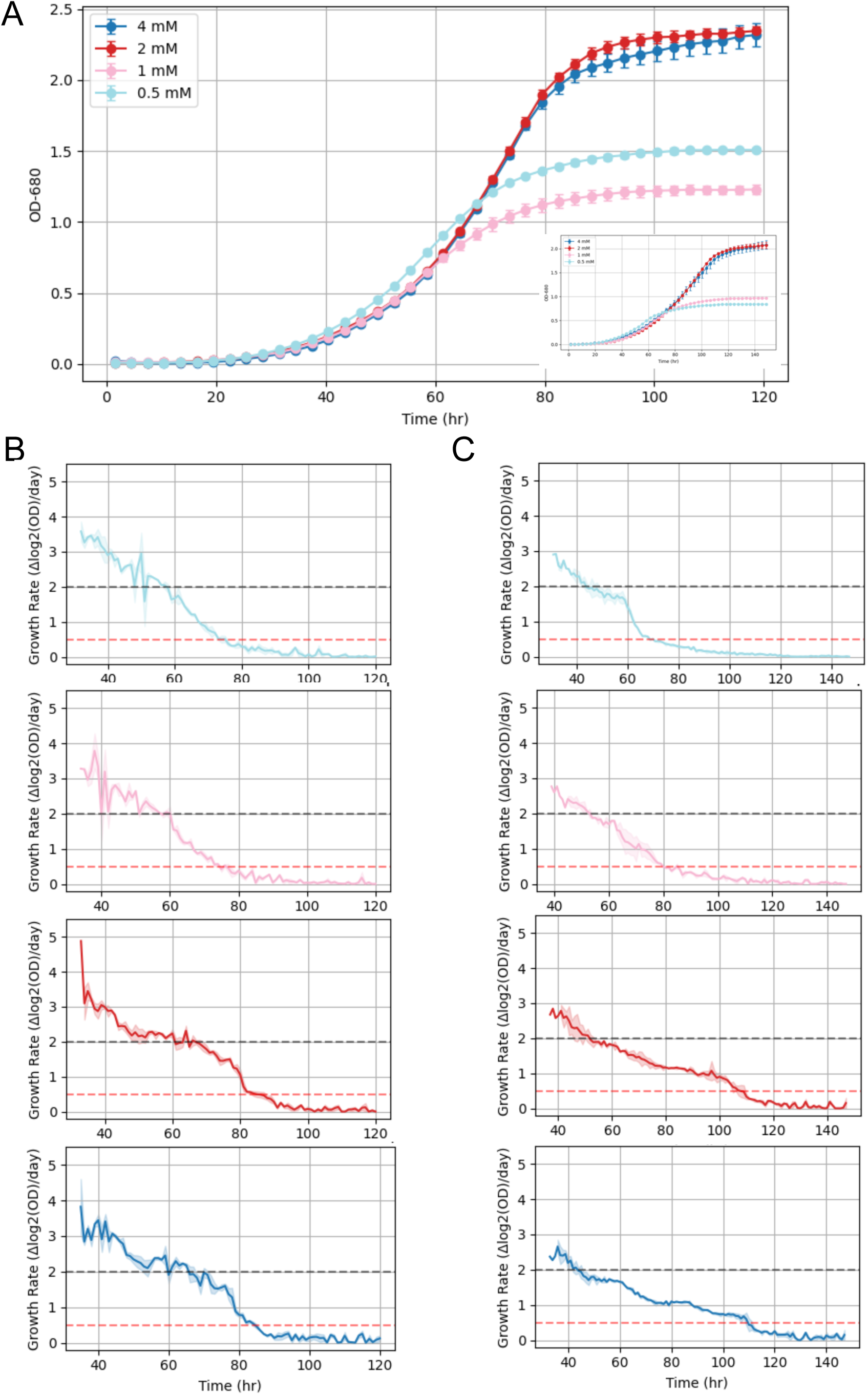
Urea-dependent growth and doubling rates. (A) Growth kinetics of *C. reinhardtii* cultured with urea as the sole nitrogen source. Strains CC-5370 (21gr, mt⁻) and CC-5119 were grown in TAP medium supplemented with 0.5–4 mM urea, and biomass accumulation was monitored by OD_680_ over time. Main panel shows growth curves for CC-5370; the inset shows corresponding data for CC-5119. (B–C) Growth rate (expressed as doublings per day) in one-hour sliding windows was calculated based on OD_680_ measurements and moving averages in one-hour sliding windows were plotted over time for each N concentration at OD_680_ ≥ 0.1 for CC-5370 (B) and for CC-5119 (C). Black dashed line indicates 2.0 as a threshold for N-replete exponential growth, and red dashed line indicates 0.5 as a threshold for the entry into stationary phase.

Under supra-saturating N conditions, NO_3_^-^- and urea-fed cultures continued slow biomass accumulation during an extended stationary phase (80–120 hours), unlike ammonium-fed cultures. Consequently, NO_3_^-^ and urea supported higher final biomass yields (OD_680_ ≈ 2.5) compared to NH_4_^+^ (OD_680_ ≈ 2.1). This extended growth suggests that NO_3_^-^ and urea cultures maintain N assimilation and anabolic activity beyond the onset of stationary phase.

### 3.2. NUE Is Maximized Under N-Limited Conditions

Using the saturation growth curves, we quantified endpoint N use efficiency (NUE), expressed as the biomass conversion rate (BCR): the OD_680_ increase per millimolar of N consumed. BCR values rose as the initial N supply decreased (Figure 4), peaking at ∼1.2 OD_680_·mM^-1^ under or below the growth saturation threshold, with comparable maxima among all N sources.

**Fig. 4.**
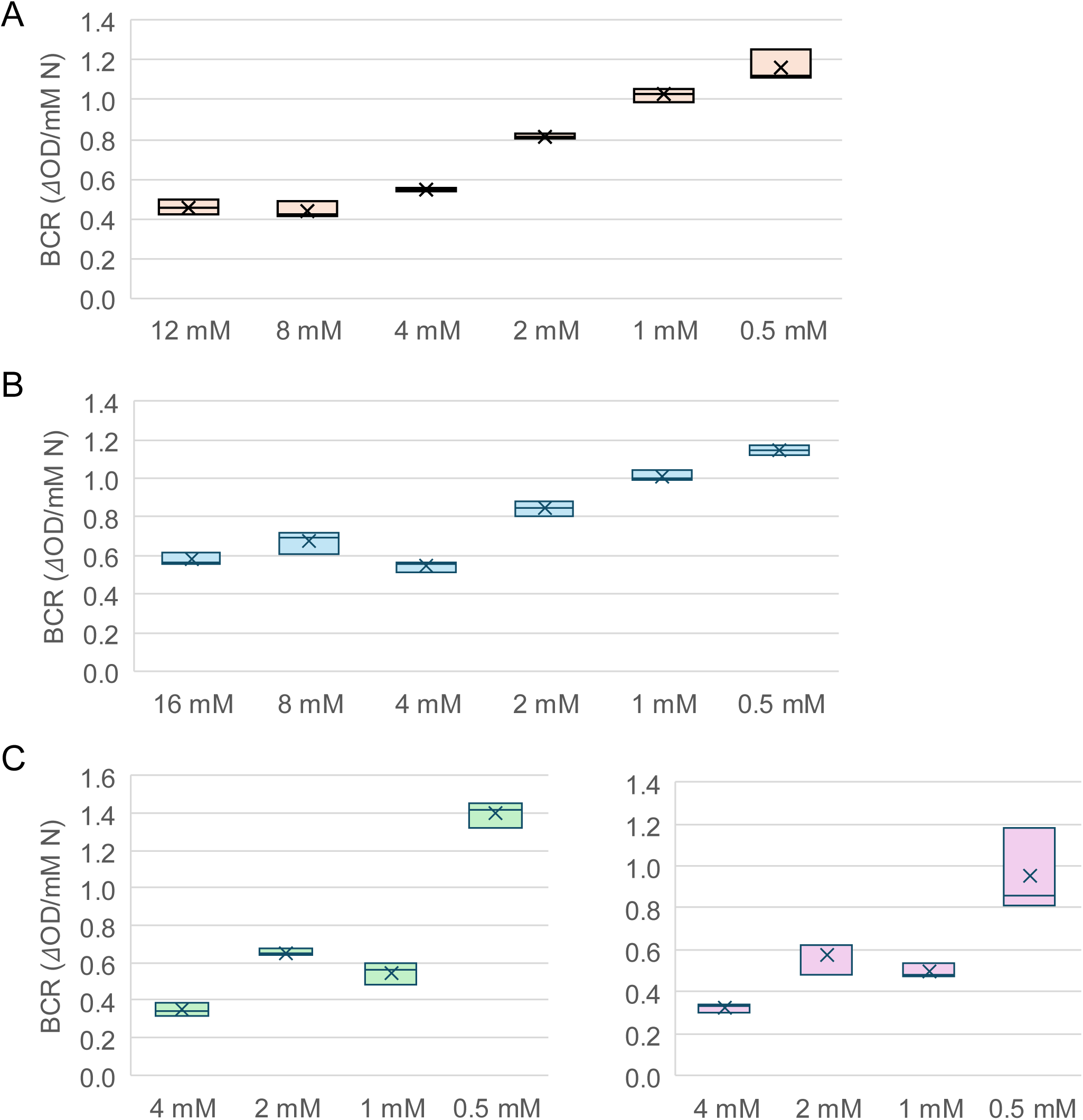
End-point biomass conversion rate (BCR) reflects nitrogen use efficiency in mixotrophic cultures. (A–C) Biomass conversion rate (BCR), calculated as final optical density (OD_680_) per mM of consumed nitrogen, was used as a proxy for N use efficiency (NUE) under mixotrophic conditions. Residual N in the medium was measured at the end of batch cultures to determine total N uptake. Boxplots represent the mean and interquartile range (25_th_-75_th_ percentile) based on biological triplicates. (A) Ammonium-fed cultures exhibited a consistent BCR of ∼0.5 OD_680_·mM^-1^ N when supplied with ≥4 mM NH_4_^+^. Below 2 mM, NUE increased, reaching up to 1.2 OD_680_·mM^-1^ N. (B) Nitrate-fed cultures showed a similar trend: BCR plateaued at ∼0.6 OD_680_·mM^-1^ N under sufficient N (≥4 mM) and increased under N limitation (≤2 mM). (C) Urea-fed cultures of both CC-5370 (left) and CC-5119 (right) followed the same pattern of increased NUE under N-limiting conditions, with BCR values rising from ∼0.4 to ∼1.2 OD_680_·mM^-1^ N as external N decreased.

At higher N concentrations, BCR declined to minima of ∼0.4 OD_680_·mM^-1^ for NH_4_^+^ and urea, and ∼0.5 OD_680_·mM^-1^ for NO_3_^-^, indicating proportionally greater N consumption per unit biomass as availability decreased. Above the threshold concentrations for NH_4_^+^ or NO_3_^-^, N consumption per biomass reached a plateau, suggesting the capacity maximum of intracellular storage, predictably constrained by the maximum N assimilation rate under specific growth conditions.

### 3.3. Differential BCR Reflects N Source-Dependent Uptake and Storage Regulation

Nitrogen uptake in *Chlamydomonas* occurs via low-affinity, high-capacity transporters or channels and high-affinity, low-capacity systems, whose expression and activity respond to external N levels (Calatrava et al., 2023). However, the interplay between uptake, biomass accumulation, and intracellular N storage under changing N availability remains less understood.

To evaluate whether distinct mechanisms are deployed under different N regimes, we quantified N consumption during the logarithmic growth phase (Figure 5). Under limiting N conditions, NH_4_^+^-fed cultures assimilated ∼1.2 mM NH_4_^+^ per OD_680_, 35% higher than the stoichiometry derived from the maximum BCR. Under excess N conditions, NH_4_^+^ uptake rose to ∼3.0 mM N per OD_680_, despite comparable growth rates (Figure 5A), implying enhanced N assimilation and accumulation beyond immediate biosynthetic demands during the logarithmic growth.

**Fig. 5.**
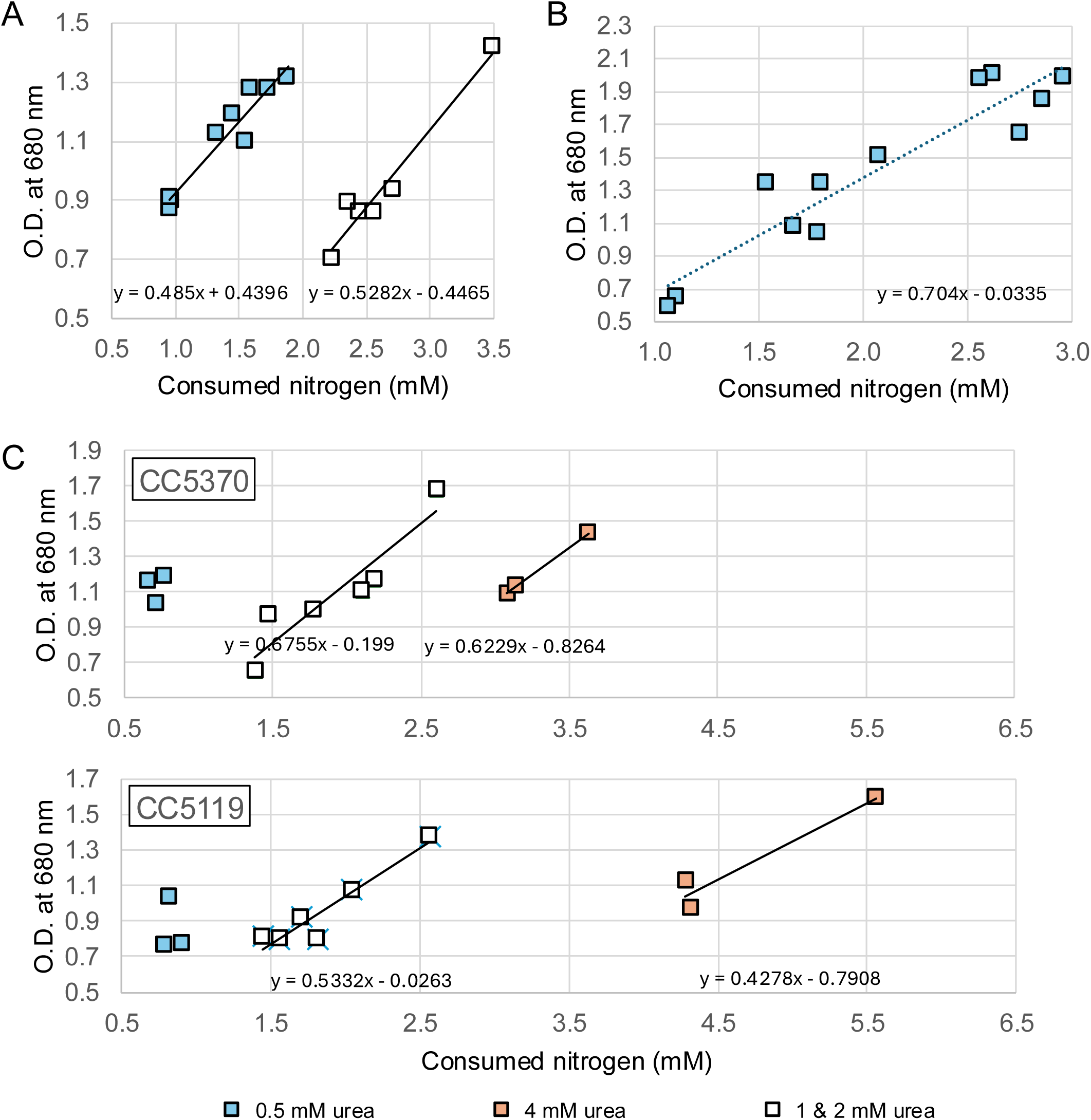
Source-dependent nitrogen uptake capacity varies with external N availability and source. (A–C) Relationships between consumed N (x-axis) and corresponding biomass accumulation (OD_680_, y-axis) during the logarithmic growth phase in *C. reinhardtii* cultures grown with ammonium, nitrate, or urea. Residual N was measured prior to the onset of stationary phase to calculate net N assimilation during exponential growth. (A) Ammonium-fed cultures (solid symbols: 1–4 mM NH_4_^+^; open symbols: 8 and 12 mM NH_4_^+^) show a nonlinear trend, with enhanced N uptake at higher external concentrations. (B) Nitrate-fed cultures (2–16 mM NO_3_^-^) display a strong linear relationship between consumed N and biomass produced, indicating consistent N uptake and assimilation efficiency across the examined concentration range. (C) Urea-fed cultures (0.5–4 mM) show a nonlinear response similar to that with ammonium, with increased N uptake at higher concentrations. The CC-5370 shows uptake capacity comparable to the highest observed in ammonium-fed culture, while CC-5119 shows further enhanced peak N uptake at 4 mM urea (orange symbol).

Urea-fed cultures showed a comparable rise in total N assimilation from 0.7 mM to 2.0 mM N per OD at 1-2 mM urea concentrations in both strains tested (CC-5370 and CC-5119), whereas two strains showed nearly 1.5 fold differences in their maximum assimilation capacities (∼3 and 4 mM per OD_680_) (Figure 5C). In contrast, NO_3_^-^-fed cultures displayed relatively constant N assimilation rates of 1.4 mM N per OD_680_ across 2–16 mM NO_3_^-^ (Figure 5B).

Together, these results indicate that N assimilation and storage are source- and availability-dependent. NH_4_^+^ and urea trigger adaptive sequestration mechanisms under high N conditions, whereas NO_3_^-^ assimilation remains steady, suggesting a less flexible uptake system under mixotrophic growth.

### 3.4. Aggressive N Uptake Supports Intracellular N Storage

To further assess whether cells actively sequester N under high availability, we conducted short-term N pulse experiments. High-density cultures were diluted into fresh medium containing 8 mM N, and N depletion was monitored over 10 and 24 hours for NH_4_^+^- and NO_3_^-^-fed conditions, respectively. During these pulses, cultures exhibited low apparent BCRs, 0.24 for NH₄⁺ and 0.13 for NO_3_^-^ (Table 1), far below the 1.2 OD_680_·mM^-1^ maximum, implying substantial N assimilation uncoupled from biomass gain.

**Table 1.** Biomass Conversion Rate (BCR) Following Nitrogen Boost in High-Density Cultures. (1) 10 hours post-harvest for NH4+ boost. (2) 24 hours post-harvest for NO3-boost. Values represent mean ± SE from biological triplicates.

| | OD<sub>680</sub> | | Residual N (mM) | | BCR<br>($\Delta\text{OD}_{680} \cdot \text{mM}^{-1} \text{ N}$) |
| --- | --- | --- | --- | --- | --- |
|  | before | after | before | after |  |
| (1) 8 mM $\text{NH}_4^+$ | 1.58 ± 0.01 | 1.92 ± 0.02 | 4.67 ± 0.18 | 3.25 ± 0.24 | 0.24 ± 0.01 |
| (2) 8 mM $\text{NO}_3^-$ | 1.44 ± 0.05 | 2.05 ± 0.13 | 6.89 ± 0.04 | 2.19 ± 0.30 | 0.13 ± 0.01 |

Estimated excess assimilation reached 2.9 mM and 6.5 mM N per OD_680_ biomass for NH₄⁺ and nitrate cultures, respectively.

If some assimilated N were distributed among pre-existing cells rather than contributing entirely to new biomass, the adjusted BCRs would be 0.38 (NH_4_^+^) and 0.45 (NO_3_^-^), consistent with the lowest steady-state BCRs observed. These findings support a model in which rapid N uptake feeds into storage reserves rather than direct growth.

### 3.5. Intracellular N Reserves Support Growth Without External Supply

To determine whether stored N directly supports subsequent biomass production, we conducted run-off bioreactor experiments in which harvested cells were resuspended in N-free medium and monitored for extended growth. Stationary-phase cultures that had depleted external N continued to increase biomass by ∼15% over 24– 48 hours, regardless of N source (Table 2A), demonstrating that internal N stores can transiently sustain growth. When logarithmic-phase cultures grown in 8 mM NH_4_^+^ received an additional 8 mM NH_4_^+^ pulse for 24 hours before resuspension (retaining 2– 3 mM residual NH_4_^+^), post-transfer growth increased by 21–23% (Table 2B), consistent with utilization of enhanced N reserves.

**Table 2.** Remobilization of N storage for growth. To test nitrogen remobilization capacity, Chlamydomonas cultures were transferred into fresh medium lacking nitrogen and their biomass increase over 48 hours was monitored to estimate the extent of internal N storage utilization. Values represent mean ± SE from biological triplicates. (A) Six-day old stationary phase cultures that had exhausted N source in the medium (below 0.05 mM). (B) Log-phase cultures at 24 hr following 8 mM N-boost (residual 1.5 ∼ 2.0 mM NH4 in the medium at the time of harvest).

| (A) |  | OD<sub>680</sub> |  |  | Remobilizable |
| --- | --- | --- | --- | --- | --- |
| Preculture | Stationary | Day 0 | Day 1 | Day 2 | N storage* |
| 4 mM NH<sub>4</sub><sup>+</sup> | 1.97 ± 0.04 | 1.39 ± 0.02 | 1.56 ± 0.01 | 1.58 ± 0.01 | 12.2% ± 1.0 |
| 4 mM NO<sub>3</sub><sup>-</sup> | 1.72 ± 0.03 | 1.22 ± 0.02 | 1.41 ± 0.02 | 1.43 ± 0.01 | 15.6% ± 0.2 |
| 2 mM Urea | 1.65 ± 0.02 | 1.16 ± 0.01 | 1.26 ± 0.01 | 1.34 ± 0.02 | 15.5% ± 0.3 |
| 1 mM Urea | 0.98 ± 0.07 | 0.69 ± 0.05 | 0.70 ± 0.04 | 0.77 ± 0.03 | 14.8% ± 3.6 |

| (B) |  | OD<sub>680</sub> |  |  | Remobilizable |
| --- | --- | --- | --- | --- | --- |
| Preculture | Strain | Day 0 | Day 1 | Day 2 | N storage* |
| 8 mM NH<sub>4</sub><sup>+</sup> | CC-5370 | 1.18 ± 0.04 | 1.37 ± 0.04 | 1.42 ± 0.04 | 21% ± 0.4 |
|  | CC-5119 | 1.62 ± 0.04 | 1.94 ± 0.02 | 1.99 ± 0.04 | 23% ± 0.6 |

In all run-off conditions, OD_720_ increased more than OD_680_, suggesting lower chlorophyll accumulation relative to total biomass compared to nutrient-sufficient cultures. This underrepresentation of chlorophyll absorbance likely reflects an N-limited physiological state, implying that stored N sustains biomass expansion even as chlorophyll biosynthesis remains constrained. Given the underestimation of biomass by OD_680_, the post-transfer biomass gain was predictably greater than 23%.

### 3.6. N Source–Dependent Regulation of Transport and Assimilation Pathways

To contextualize these metabolic flow of N sources in culture, we generated transcriptomes from NO_3_^-^-fed cultures and analyze them together with the transcriptomes from Urea-fed cultures published recently (Liang, 2025) to assess how *Chlamydomonas* regulates N uptake, assimilation, and storage-related genes under different N sources. Because N starvation broadly activates scavenging responses, expression profiles under NH_4_^+^-, NO_3_^-^-, and urea-fed conditions were compared to those from N-starved cells (Table 3).

**Table 3.**
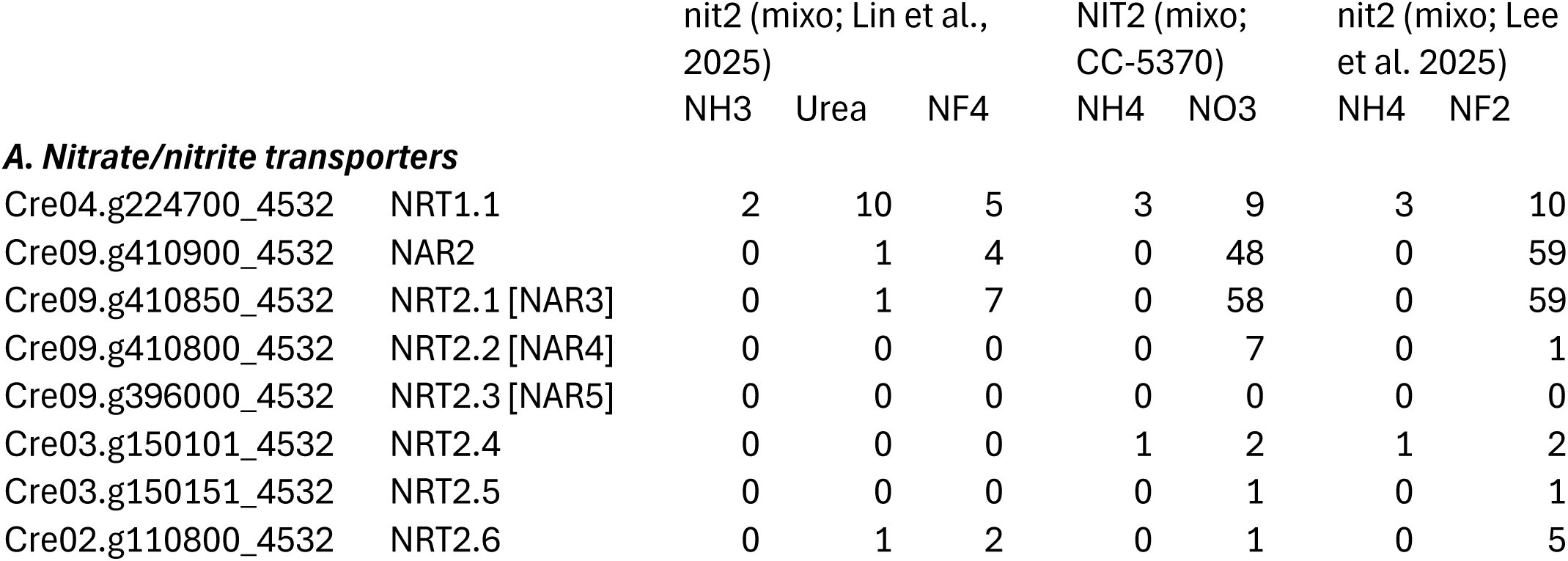
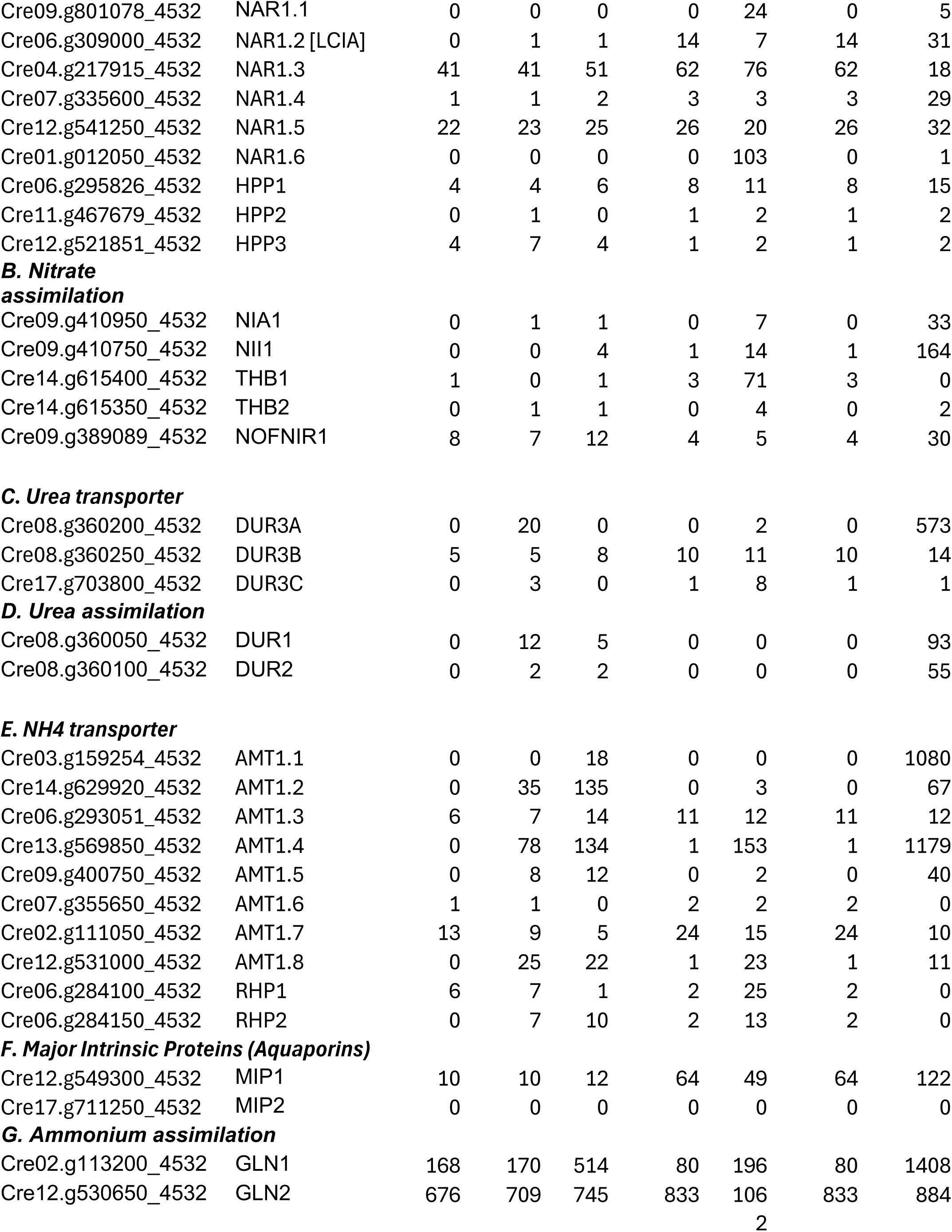
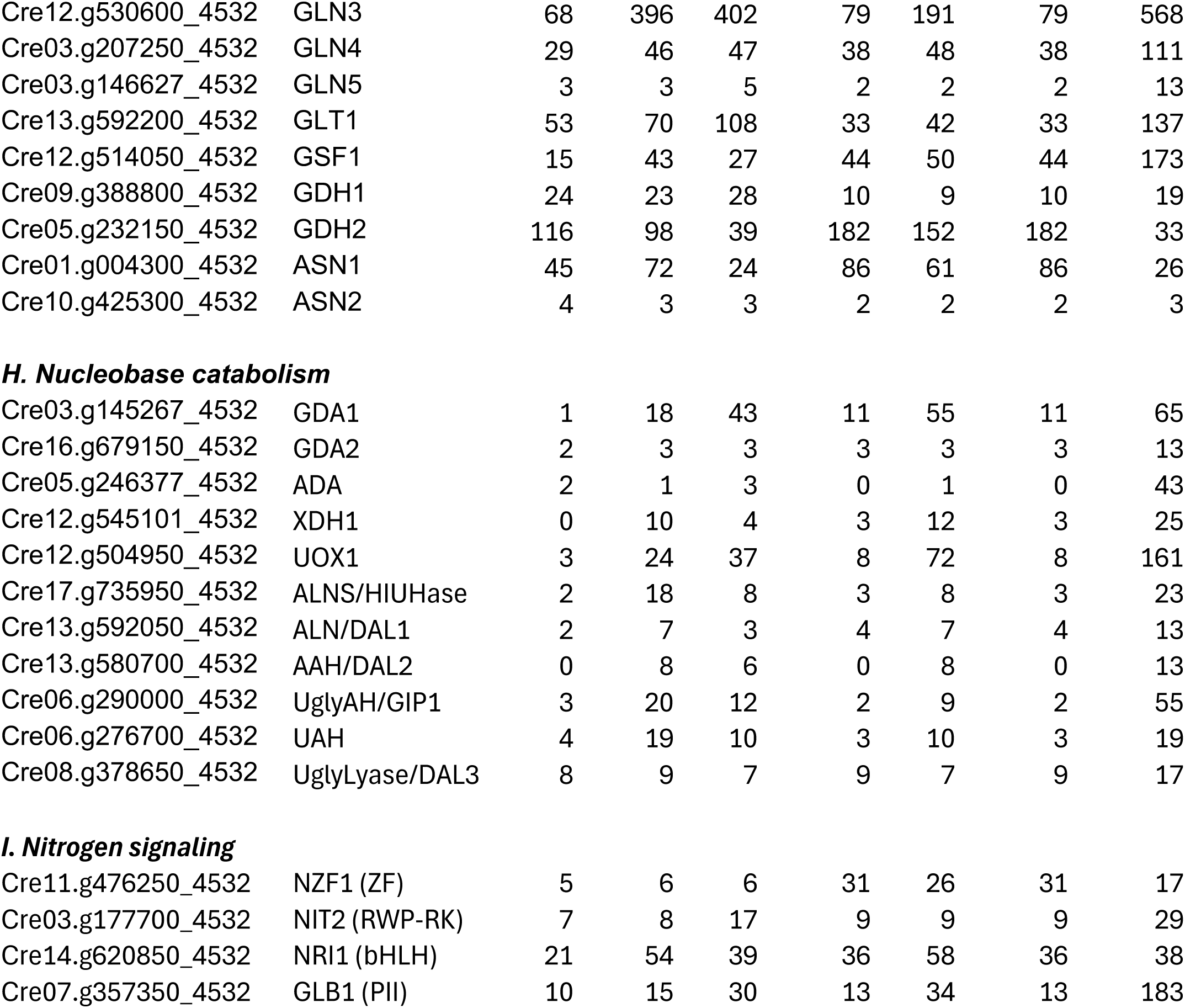
Expression profiles of N-assimilation genes in ammonium-fed, nitrate-fed, and urea-fed mixotrophic cultures. FPKM values are provided. NF4, 4 hours in N-free TAP; NF2, 2 hours in N-free TAP.

#### 3.6.1. Nitrate uptake and metabolism

Nitrate transport in NO_3_^-^-fed cultures is primarily mediated by the NRT2.1 and NRT2.2 high-affinity transporters, together with their cofactor NAR2. Strong induction of NRT2.1 and NAR2 during N starvation confirms their role in scavenging. In contrast, NAR2-independent half-size transporters (NRT2.3-NRT2.6) showed minimal expression under NO_3_^-^-fed conditions.

In land plants, anion channels are also involved in NO_3_^-^ fluxes, but yet to be characterized in single-celled algae. A potential LATS component is NRT1, a single member of the NRT1/NPF (nitrate transporter 1/nitrate peptide family) family, expanded greatly in plants, including CHL1.1, NO_3_^-^ sensor/transporter on the plasma membrane, although NRT1/MPF members are known for their broad substrate specificity.

*Chlamydomonas* has a single NRT1 family gene, NRT1.1. Low but enriched expression of NRT1 in NO_3_^-^-fed culture suggests its involvement in NO_3_^-^ uptake.

Within the FNT (formate nitrite transporter) family, NAR1.1 and NAR1.6 displayed NO_3_^-^-exclusive expression, supporting a specific role in nitrite import during NO_3_^-^ assimilation. Expression of HPP2, a homolog to the chloroplast-localized nitrite transporter characterized in plants (Maeda et al., 2014), also followed NO_3_^-^-dependent activation, suggesting functional conservation. NO_3_^-^ reduction to NH_4_^+^ proceeds through NIA1 and NII1, both upregulated under NO_3_^-^ and N-starvation. The NO_3_^-^-exclusive expression of THB1 and THB2, known NO detoxifiers, supports their function in recycling NO species during high NO_3_^-^ assimilation fluxes.

#### 3.6.2. Urea transport and metabolism

Among three urea transporter genes, DUR3A and DUR3C showed urea-exclusive expression, as previously described (Liang, 2025), while DUR3A seems to play the major route based on high expression level (20 FPKM compared to 3 FPKM of DUR3C). In *Chlamydomonas*, urea catabolism proceeds via the two-step amidolyase pathway (DUR1 and DUR2), distinct from the single-step urease pathway found in plants (Liang et al., 2025). Their expression was enriched in urea-fed cultures and strongly upregulated in N-starved conditions, implying that urea catabolism is integrated into a general N recovery program rather than being strictly source-dependent.

#### 3.6.3. Ammonium transport

Members of the AMT1 and RHP high-affinity NH_4_^+^ transporter families showed low expression under NH_4_^+^ -replete conditions, except for AMT1.7, predicted to localize to mitochondria. This supports earlier findings that high-affinity NH_4_^+^ transport systems are not detectable under N sufficiency (Ermilova et al., 2010), while low-affinity bulk transport predominates, potentially via aquaporins known for their ammonia-conducting roles in plants and animals (Loque, 2005; Kirscht et al., 2016). Among the two aquaporins in *Chlamydomonas*, MIP1 serves as a major water conducting channel in contractile vacuole that pumps out excess water from the cytosol (Komsic-Buchmann et al., 2014), whereas MIP2 expression is negligible in our transcriptomes, leaving no candidate for NH_4_^+^-LATS.

#### 3.6.4. Ammonium assimilation and storage-related enzymes

In contrast to the NO_3_^-^ and urea catabolism genes, the major N assimilation enzymes of the GS/GOGAT cycle, as well as secondary NH_4_^+^-assimilating enzymes (GDHs and ASNs), showed consistent expression across N sources. This supports earlier observations that most external N is assimilated through the chloroplastic GS/GOGAT pathway (Peltier et al., 1980). GLN3, encoding a chloroplast-localized glutamine synthetase (GS), was notably downregulated in NH_4_^+^-fed cultures, mirroring the suppression of chloroplastic AMT1.4, suggesting a coordinated regulatory response to high intracellular NH_4_^+^ availability.

Predicted guanine catabolism genes were expressed at relatively low levels in NH_4_^+^-fed cultures but were elevated in NO_3_^-^- and especially urea-fed cultures. This inverse pattern aligns with the microscopy data, where guanine crystals were abundant under NH_4_^+^ but scarce under NO_3_^-^ and urea, suggesting that urea-fed cells maintain more active guanine turnover pathway that can produce urea.

## 4. Discussion

This study reveals that *C. reinhardtii* possesses remarkably flexible N use strategies (NUS), allowing dynamic adaptation to the form and availability of N. Such plasticity reflects the organism’s evolutionary success in fluctuating environments where N frequently oscillates between reduced and oxidized forms. Under mixotrophic conditions, NH_4_^+^ appeared as the most efficiently assimilated N source, sustaining the highest growth rates, up to four doublings per day, and supporting significant intracellular N storage. In contrast, NO_3_^-^ and urea supported slower growth, consistent with their higher energetic costs and more complex enzymatic assimilation pathways. Nevertheless, comparable N use efficiency (NUE) across N sources suggests that *Chlamydomonas* can balance energetic and metabolic demands to maintain productivity under diverse nutrient regimes.

### 4.1. Source-specific N Use Strategies

A striking observation was the extended growth of NO_3_^-^- and urea-fed cultures beyond the apparent stationary phase (OD_680_ ∼2.1), eventually reaching OD_680_ ∼2.5 (Figures 1–3). This secondary growth phase, absent in NH_4_^+^-fed cultures, implies that *C. reinhardtii* cells in NO_3_^-^ or urea media may transition into a different stationary state that is also physiologically distinct from that of N-starved cells (Bogaert, 2019). Because acetate depletion marks the transition to photoautotrophic metabolism, the continued growth may indicate reactivation of photosynthetic C fixation once organic C is exhausted. This behavior aligns with earlier observations that *Chlamydomonas* readily switches trophic modes to optimize energy balance under varying nutrient and light conditions (Johnson and Alric, 2012; Bogaert et al., 2019). Future experiments manipulating CO_2_ availability and light intensity during stationary phase could directly test whether this late-phase growth indeed reflects renewed photosynthetic carbon assimilation.

In contrast, NH_4_^+^-fed cultures failed to exhibit such extended growth, even though measurable NH_4_^+^ remained in the medium. A recent study demonstrated that disruption of chloroplastic GS can alleviate NH_4_^+^-induced inhibition (Hachiya et al., 2021), implicating excess NH_4_^+^ assimilation within the chloroplast as a key trigger of metabolic stress. This finding is consistent with our observation that NH_4_^+^-fed *C. reinhardtii* cells reached a growth ceiling despite available N. It is plausible that residual NH_4_^+^ (2–3 mM) interferes with photosynthetic regulation, preventing cells from switching efficiently to phototrophic metabolism. Increasing CO_2_ supply or light intensity to promote carbon flux may relieve this inhibition by rebalancing C–N metabolism.

Alternatively, the inability to reprogram from mixotrophic to autotrophic growth could reflect regulatory inflexibility in the transition of energy metabolism. Identifying the biochemical or regulatory conditions that relieve NH_4_^+^ inhibition could therefore provide tractable strategies for mitigating NH_4_^+^ toxicity in plants and for improving crop N resilience.

Our N consumption data during logarithmic growth revealed a pronounced upregulation of N consumption per biomass in NH_4_^+^- and urea-fed cultures under N-excess conditions, suggesting that *C. reinhardtii* actively sequesters excess N into storage pools. In contrast, NO_3_^-^-fed cultures maintained a relatively constant N consumption rate during exponential growth, consistent with their highest biomass conversion rate (BCR) and indicating minimal N accumulation beyond immediate biomass needs. Interestingly, however, dense NO_3_^-^-fed cultures exhibited a transient “boost” in N assimilation upon N re-supply (Table 2B), surpassing that of NH_4_^+^-fed cells. This finding implies that NO_3_^-^-grown cells retain the metabolic potential to rapidly scale up N assimilation, perhaps through regulatory activation of nitrate reductase and associated C-N metabolic pathways. Elucidating the molecular basis of this assimilation boost could provide insights into optimizing N flux and NUE in both algal and crop systems under fluctuating N inputs.

### 4.2. What Determines N Assimilation Capacity?

In the short term, N assimilation capacity is determined by the combined activities of N uptake and assimilation enzymes, particularly those of the GS/GOGAT cycle. The rapid N uptake observed during boost experiments implies coordinated activation of low-affinity transport systems (LATS) and downstream assimilation capacity. While NH_4_^+^ and NO_3_^-^ LATS are well characterized in *C. reinhardtii* (Calatrava et al., 2023), urea uptake primarily depends on DUR3-type transporters (Liang et al., 2025). Under N-replete conditions, high-capacity LATS systems likely place increased demand on GS/GOGAT-mediated assimilation. Because this cycle requires a continuous C skeleton supply, the overall assimilation capacity ultimately hinges on the coordination between N metabolism and photosynthetic or acetate-derived C flow.

Our use of mixotrophic growth minimized C limitation and allowed the full expression of N assimilation capacity. Previous studies have shown that *C. reinhardtii* can achieve comparable growth rates under photoautotrophic conditions when supplied with 5% CO_2_ (Munz et al., 2020), indicating that its photosynthetic output can sustain high rates of inorganic N assimilation. It remains to be determined whether N consumption and storage during photoautotrophic growth parallel those observed under acetate-fed mixotrophy. If photosynthetic C fixation cannot fully support the high N assimilation demands characteristic of mixotrophic growth, the degree of excess N storage during photoautotrophy could serve as a quantitative readout of N-C coordinated assimilation capacity. Such an approach would provide a practical framework to evaluate strategies aimed at improving C-N coordination and enhancing NUE in photosynthetic production systems. Ultimately, engineering stronger coupling between GS/GOGAT flux and photosynthetic energy supply may yield crops with both high biomass productivity and improved N economy.

### 4.3. N Storage Solutions Integrated to NUS

The contrasting N storage behavior among nitrate-, NH_4_^+^-, and urea-fed cells suggests that *Chlamydomonas* allocates N differently depending on its biochemical form and growth phase. In the short term, surplus N may be incorporated into ribosomes and photosynthetic enzymes such as Rubisco, effectively increasing the cell’s biosynthetic capacity. Because *C. reinhardtii* cells can increase their biomass more than tenfold within a single cell cycle (Cross and Umen, 2015), such N incorporation directly supports biomass production of daughter cells rather than long-term storage in dividing cells. Dividing nitrate-fed cells appear to channel assimilated N primarily into rapidly turn-over pools rather than into long-term reserves. This would explain the consistently high BCR with minimal N storage but enhanced assimilation under dense, slower-growing conditions (Figure 4A). Biochemical dissection of cellular N partitioning into proteins, nucleic acids, and specialized N-rich compounds (e.g., arginine, polyamines, and guanine) will be essential for defining the metabolic routes that underpin source-dependent NUS.

Microalgae-specific N storage mechanisms include guanine storage vesicles (GSVs), which accumulate crystalline guanine as a dense, space-efficient N reserve (Mojzes et al., 2020; Goodenough et al., 2025). Unlike plant cells, which can store nitrate within central vacuoles, *Chlamydomonas* lacks large vacuolar compartments, making GSVs a likely evolutionary adaptation for nitrogen storage in compact cells. Analogous to polyphosphate granules for phosphorus (Goodenough et al., 2019), guanine crystals provide a metabolically inert nitrogen sink that can be mobilized under limitation. Future work combining microscopy, isotope tracing, and mutant analyses will clarify the quantitative contribution of GSVs to N homeostasis in microalgae.

Arginine and its derivatives represent another major N sink in photosynthetic organisms (King and Gifford, 1997; Winter et al., 2015). In cyanobacteria, arginine-derived polymers such as cyanophycin serve as N reserves that can be mobilized during starvation (Forchhammer and Selim, 2019). Similarly, in plants and *C. reinhardtii*, excess ammonium rapidly elevates glutamine and arginine levels, while other amino acids remain relatively stable (Lee et al., 2012; Hachiya et al., 2021; Urra et al., 2022). Arginine surge can be attributed to the glutamine-sensing PII protein, which derepresses N-acetylglutamate kinase (NAGK), the rate-limiting enzyme in arginine synthesis (Chellamuthu et al., 2014). Arginine thus functions as an immediate sink for newly assimilated N. Whether arginine or its derived polyamines contribute to long-term N storage in microalgae remains unclear (Theiss et al., 2002; Park et al., 2015).

Notably, arginine-fed cultures exhibited transcriptional and physiological signatures typical of N-starved cells (Munz et al., 2020; Lee et al., 2025), suggesting that arginine catabolism intersects with N starvation signaling pathways (Winter et al., 2015; Liebsch et al., 2022).

The ureid biosynthesis and catabolic pathways are among the most strongly upregulated during the early hours of N starvation and represent major regulatory targets of NRI1, a bHLH transcription factor that represses N scavenging genes under N replete conditions (Schmollinger et al., 2014; Jia et al., 2022). Given that guanine storage vesicles constitute a principal N reservoir, the induction of ureid metabolism genes such as GDA1 and XDH1 likely reflects the mobilization of stored guanine into ammonium for reuse (Table 4). In concert with arginine catabolism, this integration of guanine turnover into N starvation signaling may enable cells to sustain anabolic metabolism by tapping into internal N reserves. Elucidating this regulatory link between N storage and signaling will be key to understanding how microalgae coordinate N allocation with environmental nutrient fluctuations.

## 5. Conclusions

In summary, our findings highlight the extraordinary versatility of *C. reinhardtii* in managing N resources through coordinated regulation of transport, assimilation, storage, and metabolic reprogramming. This plasticity provides a distinct ecological advantage in natural environments where N availability is highly variable. From an applied perspective, understanding these adaptive strategies offers valuable opportunities for improving N use efficiency in both algal biotechnologies and terrestrial crops. By integrating insights from *Chlamydomonas* into agricultural engineering, it may be possible to design N-smart production systems that maximize biomass yield while minimizing N input, contributing to more sustainable bioresource and food production.

## Highlights

- *C. reinhardtii* achieves comparable final biomass across ammonium, nitrate, and urea when provided at saturating concentrations, but exhibits the highest growth rate with ammonium, reaching to **4 doublings per day** under mixotrophic conditions.
- Under nitrogen-limited conditions, the cells convert available nitrogen into biomass at an efficiency of approximately **1.2 OD/mM N**, establishing a baseline nitrogen use efficiency (NUE) for the wild-type strain.
- **Intracellular nitrogen storage is substantial** and varies with both nitrogen source and growth phase, enabling continued growth during periods of transient nitrogen deprivation through rapid assimilation when nitrogen is abundant.
- Transcriptomic data support **a highly flexible nitrogen response program**, with differential regulation of uptake and assimilation genes tailored to the nitrogen source.
- Together, these findings deepen our understanding of **nitrogen use strategies** in microalgae and underscore the utility of *C. reinhardtii* as a model system for engineering nitrogen-efficient crop and algal production systems.

## ACKNOWLEGEMENTS

(A) J. Lee supported by Discovery Grants 418471–12 from the Natural Sciences and Engineering Research Council (NSERC) and by University of Manitoba internal funds 325895; W. Mallawarachchi, M. Ziaeian, and H.M. Dharmasiri by University of Manitoba GETS and SEGS support; M. Tibule by University of Manitoba Summer Reseach support. *Chlamydomonas* strains for this research were obtained from the Chlamydomonas Resource Center, funded by the US National Science Foundation.

## Abbreviations

N: nitrogen
C: carbon
NUS: nitrogen use strategies
NUE: nitrogen use efficiency
GS/GOGAT: glutamine synthetase/glutamate synthase
LATS: low-affinity transport system
HATS: high-affinity transport system
NAGK: N-acetylglutamate kinase
GSV: guanine storage vesicle
FPKM: fragments per kilobase transcripts per million reads

